# Effect of time-series length and resolution on abundance- and trait-based early warning signals of population declines

**DOI:** 10.1101/568600

**Authors:** A.A. Arkilanian, C.F. Clements, A. Ozgul, G. Baruah

**Affiliations:** Department of Biology, McGill University, Montreal, Quebec, H3A 1B1, Canada; Department of Evolutionary Biology and Environmental studies, University of Zurich, Winterthurerstrasse 30, 4055 Zurich; School of Biosciences, University of Melbourne, Parkville VIC 3052, Melbourne, Australia

**Author notes:** Corresponding author: Gaurav Baruah.

**Keywords:** early warning signals, population collapse, sampling, time-series length, trait-based EWS

## Abstract

Natural populations are increasingly threatened with collapse at the hands of anthropogenic effects. Predicting population collapse with the help of generic early warning signals (EWS) may provide a prospective tool for identifying species or populations at highest risk. However, pattern-to-process methods such as EWS have a multitude of challenges to overcome to be useful, including the low signal to noise ratio of ecological systems and the need for high quality time-series data. The inclusion of trait dynamics with EWS has been proposed as a more robust tool to predict population collapse. However, the length and resolution of available time series are highly variable from one system to another, especially when generation time is considered. As yet it remains unknown how this variability with regards to generation time will alter the efficacy of EWS. Here we take both a simulation- and experimental-based approach to assess the impacts of relative time-series length and resolution on the forecasting ability of EWS. We show that EWS’ performance decreases with decreasing length and resolution. Our simulations suggest a relative time-series length between ten and five generations and a resolution of half a generation are the minimum requirements for accurate forecasting by abundance-based EWS. However, when trait information is included alongside abundance-based EWS, we find positive signals at lengths and resolutions half of what was required without them. We suggest that, in systems where specific traits are known to affect demography, trait data should be monitored and included alongside abundance data to improve forecasting reliability.

## Introduction

Anthropogenic pressures have long been known to reduce the resilience of ecological systems, leaving them vulnerable to transitioning into undesirable states where the systems’ ability to provide valuable ecosystem services is diminished. Such undesirable transitions have occurred in multiple systems such as in the global whale stock collapse of the 20th century due to overfishing (Hilborn et al. 2003, Clements et al. 2017); lake eutrophication through heavy nutrient input (Smith and Schindler 2009); or coral bleaching as a result of increased ocean warming (Hughes et al. 2017). In many cases recovery from such a perturbed state can be difficult as complex systems such as those seen in ecology often show hysteresis (Folke et al. 2004, Scheffer et al. 2009), thus driving a need to minimize impacts on biological systems, as well as a developing effective methods to monitor them (Costanza et al. 1997).

Early warning signals (EWS) have been shown to predict population collapses (Wissel 1984, Drake and Griffen 2010, Dai et al. 2012, Clements and Ozgul 2018) and shifts in ecosystem states (Scheffer et al. 2009, Carpenter et al. 2011). These indicators provide the possibility to intervene and reverse these undesirable events (Biggs et al. 2009, Pace et al. 2017). Classical EWS are statistical signatures which arise as a result of a phenomenon known as critical slowing down (CSD) that occurs prior to an ecosystem transition (Dakos et al. 2008, Scheffer et al. 2009, Clements and Ozgul 2018). CSD occurs as a system loses stability in the face of increasing external stress and takes longer to return to its original equilibrium state (Wissel 1984). Directly measuring CSD requires monitoring the return rate of the system which is challenging to do in natural systems. Alternatively, whether a system is experiencing CSD can be inferred through other statistical metrics measured over state-based data, for example abundance time series. Some proposed statistical signatures related to the return rate of a dynamical system are variance and autocorrelation, although various other metrics have also been developed (Dakos et al. 2012a). Increases in variance and autocorrelation in abundance time series, known as classical abundance-based EWS (Drake and Griffen 2010, Dai et al. 2012), are shown both theoretically and experimentally to occur in systems approaching transition (Drake and Griffen 2010, Carpenter et al. 2011, Dakos et al. 2012b, Clements and Ozgul 2016b).

Increases in variance and autocorrelation, along with other classical abundance-based EWS, provide an ideal generic method that responds to the dynamics of the system independent of any system-specific data such as intrinsic demographic processes and extrinsic environmental factors. However, the performance of these classical EWS has been questioned in numerous simulations (Hastings and Wysham 2010, Boerlijst et al. 2013b, Clements et al. 2015, Burthe et al. 2016, Clements and Ozgul 2016a, Dutta et al. 2018) as well as in experimental and field data (Wilkinson et al. 2017, Pace et al. 2017). Recent studies on data quality have shown that these signals might require high-resolution time-series data to produce reliable forecasting (Clements et al. 2015). Given that EWS are statistics derived from abundance time series, the quality of data available is critical to obtain a strong forecast by EWS and temporal limitations might have many consequences.

In monitoring programs from natural ecological systems high resolution data can be hard to achieve. In addition, data can be spatially and temporally limited due to constraints on resources. Thus, data from ecological systems can often present with short time-series lengths, low sampling resolutions, or both (Clements et al. 2015). Further, abundance time-series data collected in field or experimental populations can vary greatly in temporal quality. For instance, in a laboratory experiment, populations of *Didinium nasutum* were sampled roughly once every two generations for 45 days before a collapse of the population occurred (Clements and Ozgul 2016a) while in a field experiment data on a lake system was collected once a day during the summer season for three years (Carpenter et al. 2011).

Similarly, in wild populations demographic data is generally collected monthly (van Benthem et al. 2017) or annually (Walle et al. 2018). Due to these differences in sampling effort the temporal quality of the time series relative to the process rate of the system (for example the generation time of the species sampled) will vary (Clements and Ozgul 2018). Previous work suggested that the length and resolution of data being analyzed in relation to the process rate, that is: the relative length and resolution, could alter the rate at which a tipping point occurs (Spanbauer et al. 2016). Following this, it has been shown that the speed at which a tipping point occurs can affect the detectability of abundance-based EWS (Clements and Ozgul 2016b). It is thus unknown how the temporal quality of the time series relative to the generation time of the organisms being monitored will affect the detectability of abundance-based EWS. Given that wild populations are monitored using varied sampling efforts, it is important to further our understanding of the relative length and resolution of time series needed to derive reliable EWS. In turn, this would help lay the foundation for generalizable guidelines for the monitoring of populations with the aim of reliably predicting population declines, or whether time series that are currently available will be suitable for detecting abundance-based EWS.

Composite EWS have been proposed as a more reliable method whereby multiple leading indicators are combined to increase overall forecasting ability (Drake and Griffen 2010). Recent work has used this composite approach to drive the inclusion of fitness related phenotypic trait data, specifically body size, alongside abundance-based methods to create trait-based EWS (Clements and Ozgul 2016a, Clements et al. 2017). The motivation behind the use of fitness-related trait data combined with leading indicators comes from a body of work providing evidence that individual traits affected by changes in the external environment are linked with concurrent demographic changes (Ozgul et al. 2009, Pigeon et al. 2017, Baruah et al. 2019b). In particular, individual plasticity in body size has been shown to buffer environmental change to higher trophic levels for example in the face of reduced food availability, climate change, and increased pollution (Brown et al. 2004, Cheung et al. 2013), For example, previous study has shown that shifts in body size can accompany a transition in diatom communities (Spanbauer et al. 2016). In addition, trait-based signals derived from body size data have been shown to be more robust than traditional abundance-based leading indicators (Clements and Ozgul 2016a, Clements et al. 2017, Baruah et al. 2019b). There remains a need to fully assess whether the inclusion of body size data leads to any considerable improvement in forecasting population collapses in the face of common data quality issues, such as shortened time-series lengths.

In this paper, we use model-simulations and data from microcosm populations of *Didnium nasutum* (Clements and Ozgul 2016a) to test and compare the effects of sampling length and resolution on the detectability of population collapse by both classical abundance-based EWS as well as trait-based EWS. We first investigated the strength and reliability of abundance-based EWS across a range of sampling lengths and resolutions. Subsequently, we tested whether the inclusion of trait dynamics (body size) can increase forecasting ability even when data is sparse.

## Methods

### 2.1 Simulations: transcritical and fold bifurcation model

We first modeled the logistic growth of a population that moves from an underexploited state to critically exploited state through a non-catastrophic transcritical bifurcation. The population is forced through the transcritical bifurcation via a linear harvesting regime. The dynamics of this population are given by:

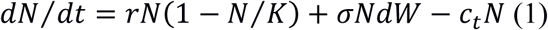

where, *r* is the growth rate of the population (0.5 individuals/day), *K* is the carrying capacity (100 individuals), *c_t_* is the harvesting rate, and *σNdW* is the Gaussian distributed white noise process with mean 0 and standard deviation *σ* (1.5). Time step *dt* used was 0.3 for each of the stochastic simulations and was implemented using the Euler approximation. The simulation was run for 100 total time steps. Next, to simulate the dynamics of population collapse via a fold catastrophe, we used a second model where harvesting of individuals of the population followed a non-linear function (May 1977, Scheffer 2009). The parameters in this model are identical to those of the transcritical catastrophe model except for the addition of *h*, the half-saturation constant:

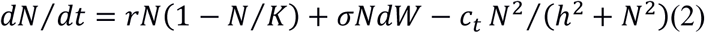

### 2.2. Population collapse and abundance-based EWS

We simulated population collapse for the two models by increasing the value of the harvest parameter ct linearly with time. We used three different levels of forcing (Clements and Ozgul 2016b) : 1) slow forcing: where the rate of forcing increased linearly from 0.03 and 0.0015 in fold and transcritical models respectively; 2) moderate forcing: where the forcing parameter *ct* increased linearly at the rate of 0.045 and 0.0025 in fold and transcritical models respectively; fast forcing: where the forcing parameter *c_t_* increased linearly at the rate of 0.07 and 0.004 in fold and transcritical bifurcation models respectively. For each simulated population’s time-series data, we estimated the bifurcation time point by fitting GAMS (Generalized Additive Modelling) to 1/*N* (*dN*/*dt*) over time *t.* The time point at which 1/*N* (*dN*/*dt*) < 0 is then our estimated bifurcation time point. For abundance-based EWS’ analyses we discarded abundance time-series data after the estimated bifurcation point. We then applied generic EWS of population collapse by using the *earlywarnings* package (Dakos et al. 2012a) in R version 3.5.2 (R Core Team 2018). Specifically, we used two early warning indicators: autocorrelation at first-lag (*ar(1)*) and standard deviation (*sd*). Other indicators such as return rate, first-order autoregressive coefficient, coefficient of variation can theoretically be derived from these two main indicators. We used Gaussian detrending to remove any trends in the abundance time-series data. To quantify the strength of population collapse, we calculated Kendall’s Tau correlation coefficients of the statistical indicators over time. Strong positive Kendall’s Tau correlation of the statistical indicators (*sd*, ar(1)) with time would indicate an approaching population collapse (Dakos et al. 2012b). We further quantified the rate of false negatives as the number of times Kendall’s Tau value calculated to be less than or equal to zero within the set of replicate time series. While previous work has suggested that a strong trend is indicated by a Kendall’s tau correlation approaching one (Dakos et al. 2012b), we use this false negative metric as an alternative visualization to presenting raw Kendall’s tau values.

### 2.3 Effect of sampling in relation to simulated population dynamics on EWS

To study the effect of varying resolutions on the detectability of collapse, we subset the data into four datasets that varied in their resolution: as the generation time for the populations in the simulation models is t=1, sampling of the abundance time series was done every quarter of *t*, every half of *t*, every *t*, and every two *t.* Interpolation between points was not performed. With these four datasets, we explored the effect of different sampling resolutions on the efficacy of EWS forecasting. Specifically, we assessed the resolution required to detect positive EWS. We assessed the effect of these sampling regimes on EWS for the three different levels of environmental forcing as mentioned in section 2.2. Next, we calculated Kendall’s tau correlation coefficients as a measure of the strength of EWS and rates of false negatives as a measure of the reliability of EWS for these sampling regimes and for each level of environmental forcing.

### 2.4 Effect of varying length of simulation abundance time series on EWS

To quantify the effect of varying lengths of abundance time series on EWS we used the quarter generation sampling resolution dataset acquired following section 2.3. We used this resolution as it allowed us to explore the largest range of time-series lengths. Next, we quantified the strength and reliability of EWS on the entire length of the abundance time series and then re-quantified it using the same time series but with the earliest data point removed (or the furthest data point from the bifurcation point). We repeated this process by sequentially removing the earliest data point and re-performing EWS analysis until the time series was 6 data points, where it becomes too short for any meaningful analysis. *sd* and *ar(1)* that were estimated on abundance data using the ‘*earlywarnings’* package typically uses a sliding window approach where the window size is generally 50% of the time series length. At 6 data points, rolling window size of 50% of the time series would be just two data points, where estimating autocorrelation and standard deviation would lead to spurious values. We performed this length reduction analysis on time series from the three forcing scenarios mentioned in section 2.2. Finally, we calculated Kendall’s tau correlation coefficients as a measure of the strength of EWS and rates of false negatives as a measure of the reliability of EWS for these sampling regimes and for each level of environmental forcing.

The parameters used in our simulations were chosen such that in all three forcing scenarios populations persisted long enough to return sufficiently long time series. This allowed us to later reduce their length and re-analyze them giving us a range of time series lengths for each resolution and forcing scenario.

### 2.5. Experimental data

In addition to the model simulations, we also analyzed an experimental microcosm dataset. In this experiment, microcosm populations were forced to collapse by varying the rate of decline in food availability over time in four different scenarios: 1) fast decline in food availability, 2) moderate decline in food availability, 3) slow decline in food availability, 4) constant food availability as the control treatment. The microcosm populations consisted of protozoan ciliate *Didinum nasutum* feeding on *Paramecium caudatum*. This particular experiment used a total of 60 replicate populations, where 15 replicates were used per treatment. In our study, we used the microcosm data only from the three different deteriorating environments (fast, moderate and slow decline in food availability). For details of the experimental design refer to Clements and Ozgul 2016a.

#### 2.5.1 Effect of varying length of experimental time series on EWS

For each of the three experimental treatments (fast, moderate, and slow), we estimated the effect of different lengths of time series on the efficacy of EWS as was done with simulation data in section 2.4 to directly compare our simulation results with the experimental data.

#### 2.5.2 Effect of sampling in relation to microcosm population dynamics on EWS

We subset the experimental data into two sampling regimes. Since *Didinum nasutum* has a generation time of roughly 2 days (Beers 1926), sampling of the abundance time-series data was done: 1) every half a generation (everyday), 2) every generation (every 2 days). Subsampling of this kind was done for each of the three experimental treatments of the microcosm population collapse. Next, as was done with simulation data, we quantified Kendall’s tau correlation coefficients as a measure of the strength of EWS and rates of false negatives as a measure of the reliability of EWS for these sampling regimes and for each level of environmental forcing.

### 2.6 Inclusion of body size data: trait-based EWS

We wanted to assess whether including trait dynamic information (body size) alongside abundance-based EWS would improve the predictability of population collapse for the scenarios of time-series length and for the sampling resolutions in the experimental data for the three forcing experimental treatments. To evaluate the utility of trait-based EWS for the different sampling resolutions and lengths of abundance time series, data from mean body size was incorporated with the leading indicators in an additive manner (Clements and Ozgul 2016a). We z-standardized abundance-based EWS (standard deviation (*sd*), autocorrelation at-lag-1 (*ar(1)*), and body size time-series data. The length of body-size time series that was used was the same as the corresponding replicate abundance time series. Before a population collapse, *sd* and *ar(1)* are anticipated to increase linearly over time, while body-size is expected to decline as food availability in the experimental treatment decreases. As a consequence, z-standardized body-size time series were multiplied by −1 so that they could be included alongside standardized abundance-based EWS. Next, standardized abundance-based EWS were then added to standardized mean body-size time series to create trait-based EWS. Next, we evaluated two trait-based EWS metrics namely *ar(1)+mean body size, sd+mean body size*. Finally, we compared these two trait-based EWS in terms of strength and reliability of forecasting a population collapse with the abundance-based statistical EWS for both the scenarios of varying time-series length and varying sampling resolution.

## Results

### 3.1 Effect of sampling resolution on abundance-based EWS (simulated data)

Reducing the resolution of the time series used for analysis with abundance-based EWS in simulated populations did not led to significant reductions in the strength of abundance-based indicators of population collapse for slow and fast forcing levels. However, for moderate forcing levels, particularly for *sd*, decreases in the resolution of the time series generally led to decreases in the median strength of the signal of population collapse (Fig. 1 – yellow, Appendix S1:Fig. S4-6). Further, at highest resolutions in the moderate forcing levels, we see the highest Kendall’s tau values indicating that time series at this resolution return the most confident forecasts capable of passing more strict thresholds than our own at a tau value of zero. While the median strength of signals (*sd* and *ar(1)*) did not generally drop with decreasing resolution (Fig.1), the proportion of false negatives increased with decreasing resolution, particularly for slow and moderate environmental forcing (Appendix S1:Table S1). For fast environmental forcing, however, false negatives increased as resolution decreased. This particular result indicated that abundance-based EWS were unable to forecast collapse of populations as resolution decreased for fast environmental forcing (Appendix S1:Table S1). For *sd*, in slow and moderate levels of environmental forcing, rate of false negative decreased slightly as resolution increased. In general, forecasts became reliable only when there was moderate level of environmental forcing as we increase time series resolution.

**Figure 1.**
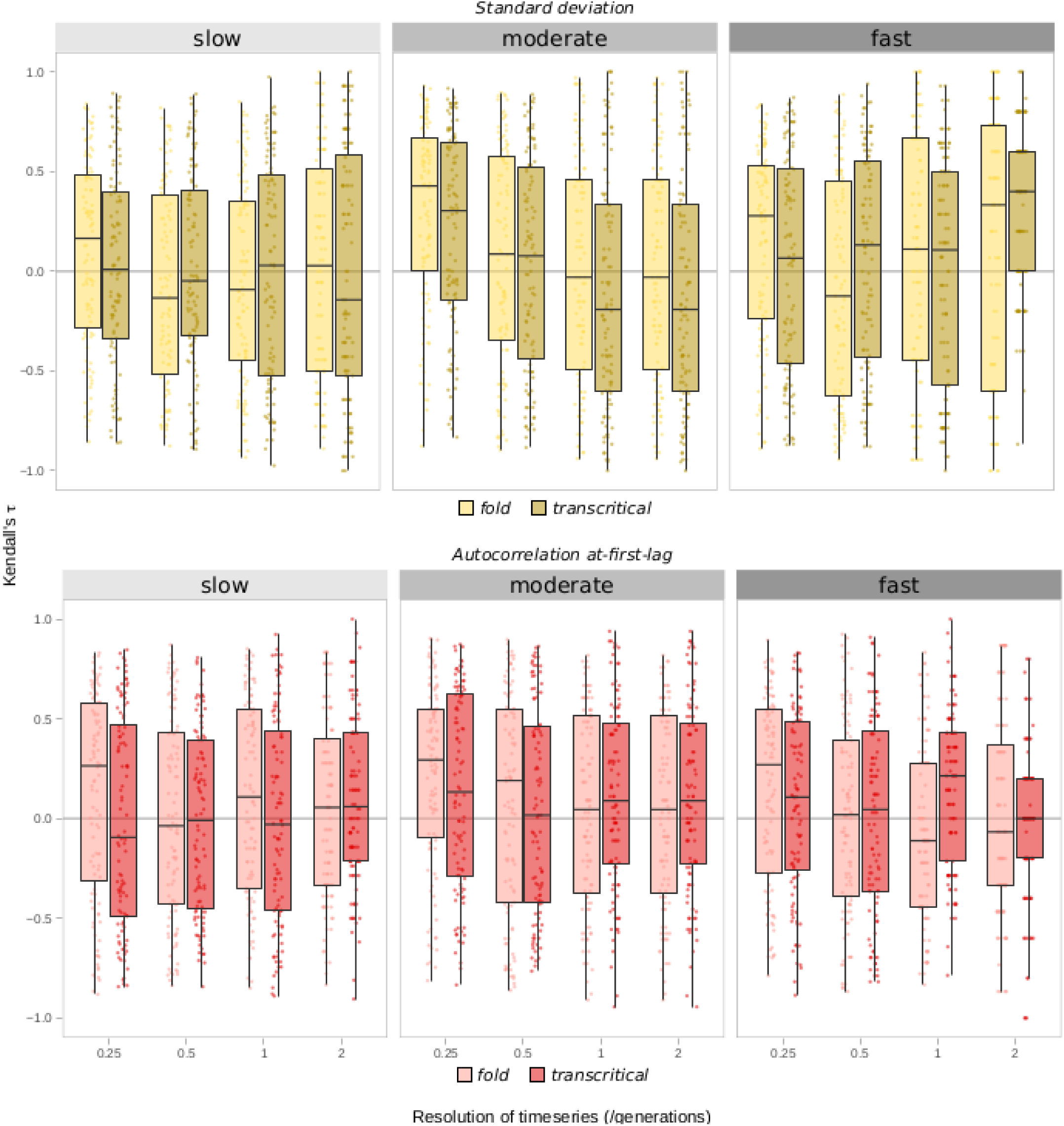
Boxplots of the strength of abundance-based early warning signals of population collapse across decreasing resolutions of sub-sampling in simulated populations subjected to collapse by harvesting. The data is split into subplots based on the bifurcation model simulated (fold, transcritical) and forcing level (slow, medium, fast). Boxplots are further split by metric used (yellow: standard deviation, red: autocorrelation at-first-lag). Each box represents the median Kendall’s tau value (shown on the y-axis) across replicate time series across three forcing level with lengths ranging from 1,000 to 40 time steps. The x-axis shows the resolution of sampling in numbers of generations such that a value of 2 indicates that sampling was performed every 2 generations and 0.25 denotes sampling every quarter generation.

### 3.2 Effect of length of time series on abundance-based EWS (simulated data)

In simulations, decreasing the length of the sampling time series before either a fold or transcritical bifurcation negatively affected the performance of abundance-based EWS *ar(1)* and *sd* across all three intensities (Fig. 2, Appendix S1:Table S2). Particularly for moderate forcing, *ar(1)* and *sd* had a strong decline in Kendall’s tau value (slope = −0.09 and R_2_ = 0.8 for *ar(1)*; slope = −0.07 R_2_=0.71 for *sd*) as length of time series decreased regardless of the type of bifurcation. Moreover, as forcing increased, the decline of signal reliability (Fig. 2B) and signal strength (Fig. 2A), as indicated by the rate of false negatives and the Kendall’s tau value respectively, saw a steeper decline. With fast forcing, when time series dropped below approximately ten generations long both EWS performed poorly with Kendall’s tau values near zero (Fig. 2A) and approximately 50% false negatives (Fig. 2B). With moderate forcing, this minimum length drops closer to 5 generation long timeseries. Finally, with slow forcing the minimum timeseries length becomes less clear, particularly for fold bifurcation, as slopes of Kendall’s tau value against length of time series were small for both *ar(1)* (slope = −0.01, R_2_ = 0.39) and *sd* (slope = −0.0003, R_2_ = −0.009). For, transcritical bifurcation and for slow forcing (dotted lines, Fig. 2A slow), however, the minimum length drops to around 5 to 10 generation long time series for both *sd* (slope = −0.075, R_2_ = 0.22) and *ar(1)* (slope = −0.09, R_2_ = 0.43).

**Figure 2.**
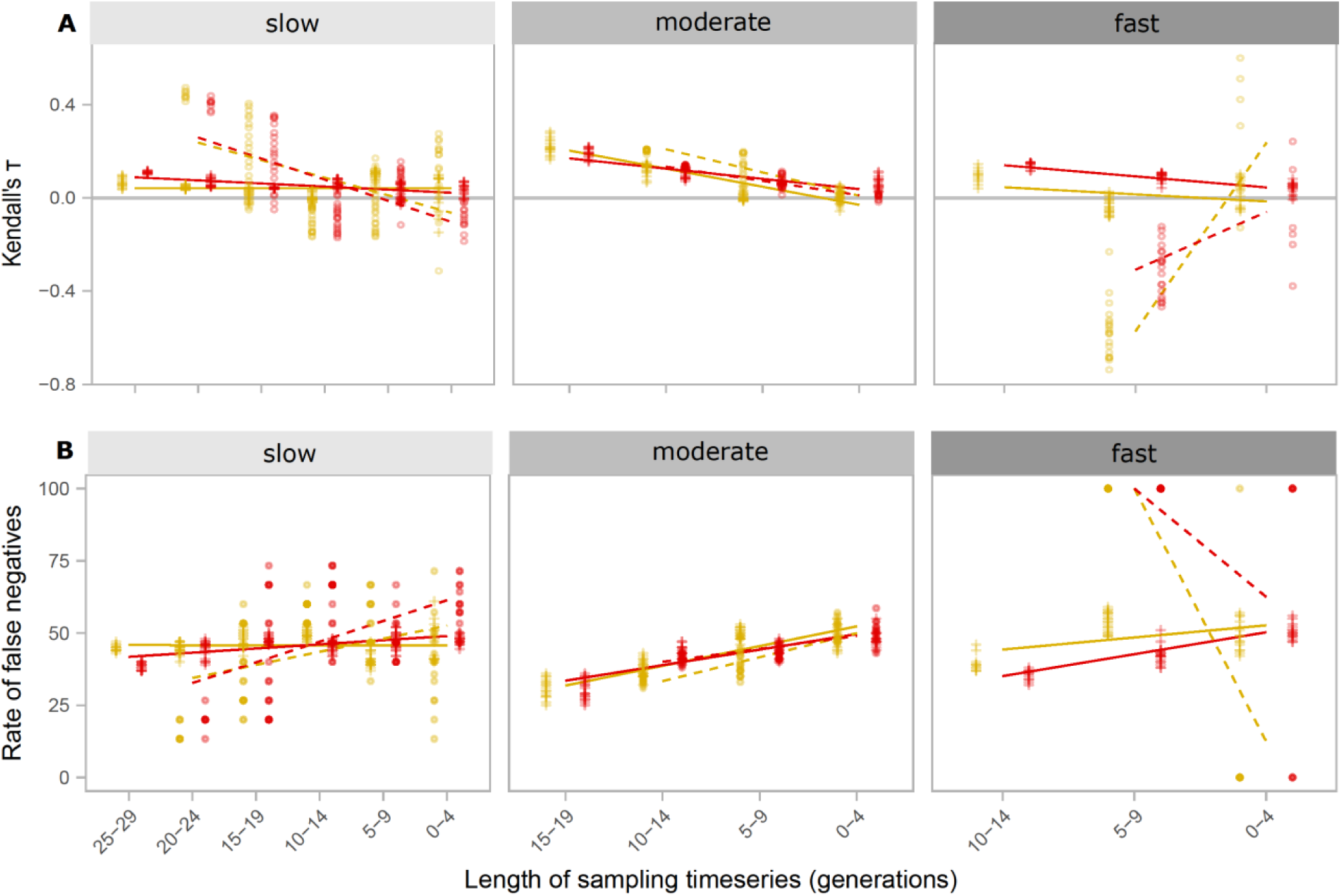
Performance of abundance-based early warning signals of population collapse across decreasing sampling time-series lengths in simulated populations subjected to collapse by harvesting. The data is split into subplots based on forcing intensity (slow, moderate, fast) and further split by metric used (yellow: standard deviation, red: autocorrelation at-lag-1) and bifurcation model simulated (solid: fold model, dashed: transcritical model). Each point represents (A) the mean Kendall’s tau correlation coefficient or (B) rate of false negatives on the y-axis of 100 replicate simulations of population collapse. X-axis represents length of time series analyzed.

### 3.3 Effect of sampling resolution on abundance and trait-based EWS (experimental data)

Decreasing the resolution of the time series in microcosm populations led to decreases in the strength of abundance-based EWS (Fig. 3), corroborating the results of model simulations (Fig. 1). The decrease in strength of was most noticeable in *sd*, while *ar(1)* remained nearly constant as resolution was decreased (Fig. 3).

**Figure 3.**
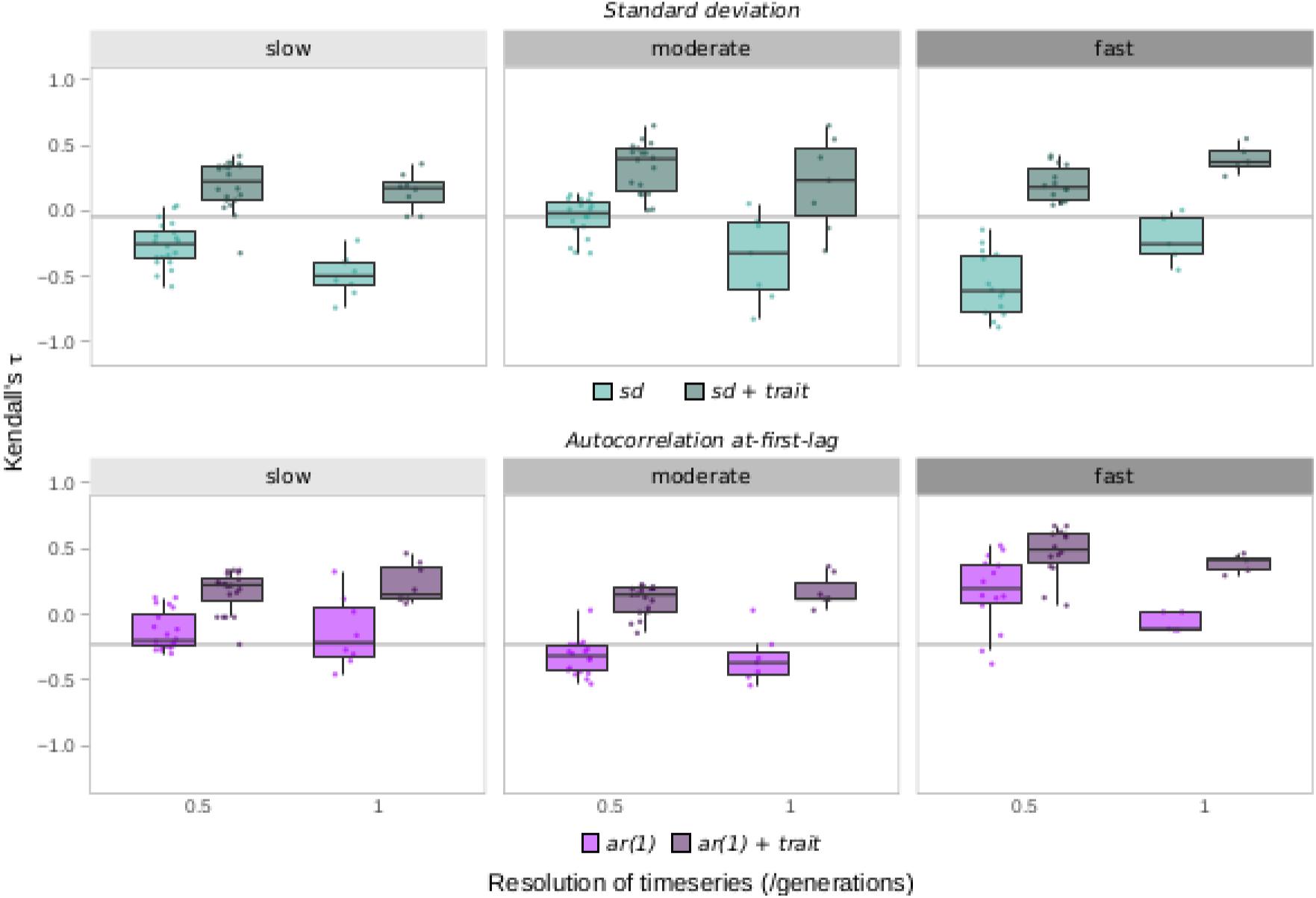
Performance of abundance-based (lighter colors) and trait-based EWS (darker colors) of population collapse across decreasing sampling time-series resolutions for the experimental data of Clements & Ozgul 2016. Plots are organized into subplots based on the forcing intensity (slow, medium, fast) and further split by metric used (blue: standard deviation, purple: autocorrelation at-first-lag). Each box represents the Kendall’s tau value (on the y axis) for each resolution studied (on the x axis). Lighter colors represent classical abundance-based EWS while the darker colours represent trait-based EWS.

Including information from mean body size data alongside abundance-based EWS to create trait-based EWS led to significant increases in strength and reliability of predicting population collapse regardless of forcing strength, or time-series resolutions (Fig. 3, dark colors).

The rate of false negatives was substantially high for abundance-based EWS across the two different time series resolution for the experimental data (appendix S1:Fig. S2). In comparison to *ar(1)*, the false negative rate of *sd* was higher in slow and fast decline in food availability across the two time series resolutions. When the resolution of time series decreased the rate of false negatives increased for abundance-based EWS, particularly for *ar(1)* (appendix S1:Fig. S2). In contrast, trait-based EWS were significantly more reliable in forecasting population collapse even when time series resolution was low.

### 3.4 Effect of length of time series on abundance and trait-based EWS (experimental data)

In agreement with the results from simulations, abundance-based EWS derived from microcosm data saw an increase in false negatives and a decrease in signal strength as the length of the sampling time series was decreased (Fig. 4A, 4B, dotted lines). This was most noticeable with fast forcing but also occurred in all forcing conditions with time series of six or less generations long. However, the drop in performance seen in microcosm data was lower than in the simulations, which is partially due to the fact that the former performs the analysis on a smaller range of time-series lengths.

**Figure 4.**
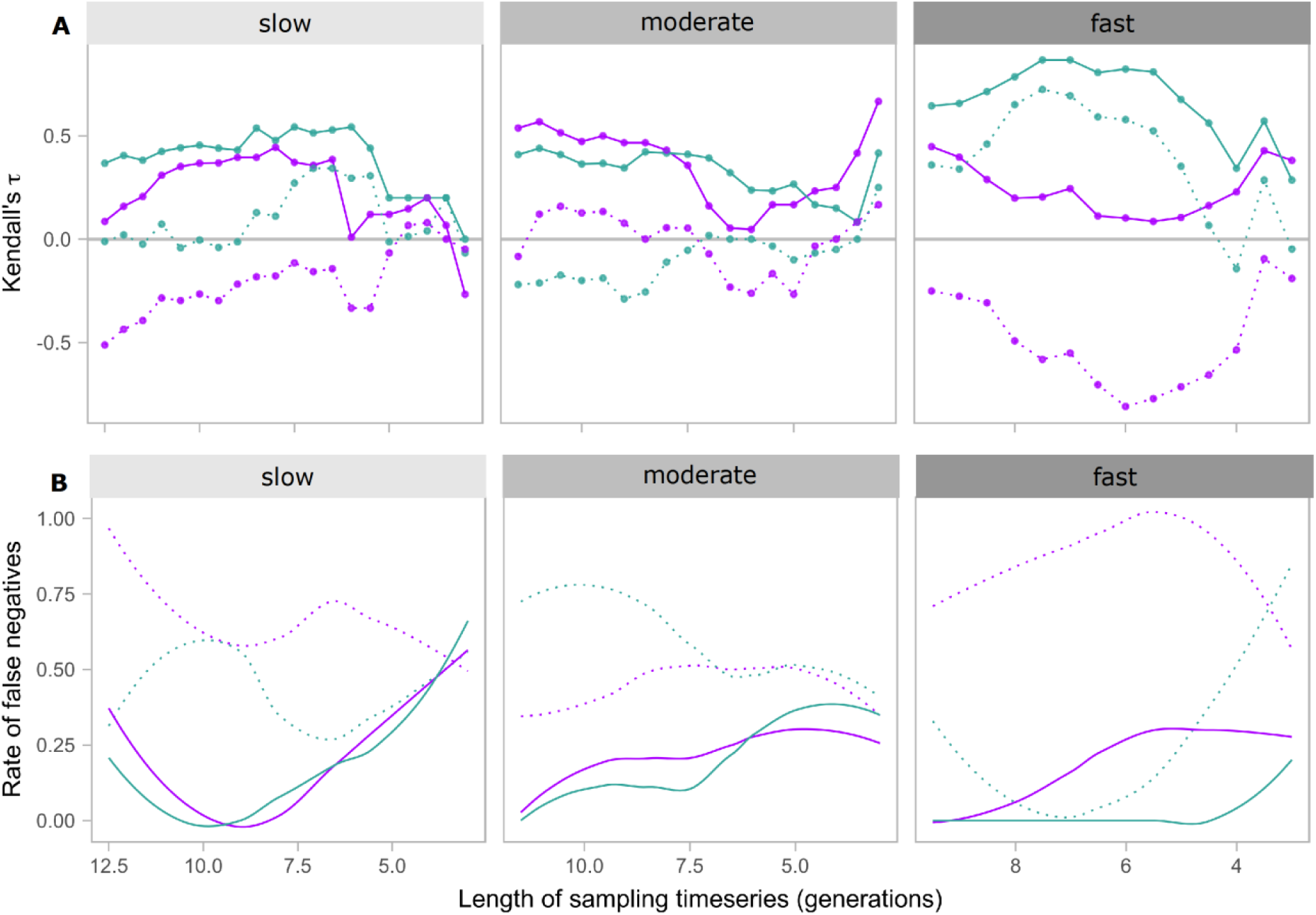
Performance of abundance-based and trait-based EWS of population collapse across decreasing sampling time-series lengths for the experimental data of Clements & Ozgul 2016. Plots are organized into subplots based on the forcing intensity (slow, medium, fast). X-axis is the length of time-series data analyzed and Y-axis denotes in (A) mean Kendall’s Tau value and in (B) rate of false negatives. Solid lines indicate EWS with body size information added (trait-based EWS) while dotted lines indicate classical abundance-based EWS. Colours indicate metric used (blue: standard deviation, purple: autocorrelation at-first-lag). False negative plots were produced using loess smoothing.

Furthermore, trait-based EWS i.e., including body size information alongside abundance EWS, significantly improved the strength as well as reliability of population collapse across different length in time-series data for different levels of environmental forcing. With trait-based EWS, Kendall’s tau value always remained positive regardless of very short time series and strength of environmental forcing (Fig 4A, solid lines). Only during slow environmental forcing scenario, and when length of time series was less than 5 generations, trait inclusive *ar(1)* was unable to predict population collapse.

## Discussion

Generic EWS would provide a unique tool for conservation prioritization and management of populations facing increased stress with changes to their abiotic environment if they are detectable prior to their collapse (Scheffer et al. 2009, Dakos et al. 2012b, Burthe et al. 2016). The attraction of generic EWS is their relative simplicity; they are easy to calculate and require only state data such as the abundance of a population (Drake and Griffen 2010, Boettiger et al. 2013, Dutta et al. 2018). Alternative approaches such as trait-based EWS have been developed with the goal of providing more reliable predictions of population collapse but require additional data to calculate (Clements and Ozgul 2016a, Clements et al. 2017). For both complementary approaches how the relative length and resolution of a time series may affect their performance has thus far remained unknown. This question is critical for our understanding of the utility and applicability of these methods (Boerlijst et al. 2013a, Boettiger and Hastings 2013). Here, using simulations and experimental data, we show that time-series length and resolution relative to the process rate of the system in question significantly influence the performance of abundance-based EWS. Further, we found that including average body size with abundance-based EWS leads to stronger and more reliable signals even when temporal length and resolution of the time series are low.

### 4.1 Effect of resolution of sampling time series on abundance-based and trait-based EWS

Reducing the resolution of time series in simulated and experimental populations subjected to different forcing levels led to a decrease in the performance of abundance-based EWS (Fig. 1 and 3). Reliability of abundance-based EWS, particularly *sd*, in forecasting population collapse for the experimental microcosms was rather poor when resolution was decreased. In fact, for the fast decline in food availability treatment false negative rate rose to 90% as time series resolution was decreased. On average, *ar(1)* performed better in forecasting population collapse than *sd* when time series resolution was manipulated for the experimental microcosm data. There were important differences in the behavior of abundance-based EWS calculated between simulated and experimental populations. Low reliability of *sd* could probably be attributed to the fact that experimental systems are inherently more stochastic. While in simulated population we found *sd* to produce the most robust signal of population collapse (Fig. 1), in experimental populations *ar(1)* performed better across varying resolutions (Fig. 3). Our simulated populations were likely subjected to smaller amounts of stochasticity in population size compared to our experimental populations. Given that *sd* is sensitive to rising stochasticity in a system (Boettiger and Hastings, 2013), one could expect *sd* to perform better when there is a high level of environmental stochasticity. Indeed, results from replicate simulations, where we varied stochasticity levels and measured the performance of abundance-based EWS suggested that *sd* performed relatively better when environmental stochasticity was higher. However, *ar(1)*, regardless of low or high environmental stochasticity, performed better than *sd.* In fact, when there was high environmental stochasticity, *ar(1)* outperformed *sd*, confirming our speculation on the performance of *ar(1)* when environmental stochasticity was high (appendix S1:Fig. S1).

In addition, abundance-based EWS showed a greater sensitivity to system stochasticity than trait-based EWS did; the latter did not present any obvious differences in performance between metrics used (Fig. 3). This is possibly an effect of adding phenotypic data to abundance data which stabilizes the signal through time and reduces stochasticity. This is an important finding if abundance-based EWS are to be used to monitor natural populations which likely have more stochasticity than laboratory populations.

As observed with abundance-based EWS, trait-based EWS also saw a decrease in their performance with decreasing sampling resolution (Fig. 3). However, these drops were in most cases very small and the performance of trait-based EWS was much better than abundance-based EWS alone. Trait-based EWS calculated with the lowest resolution in either simulated or experimental populations still derived a more reliable forecast than abundance-based EWS calculated with the highest possible resolution.

Our findings suggest that abundance-based EWS derived from autocorrelation at-first-lag would perform better in natural populations subjected to high levels of stochasticity when compared with standard deviation derived signals when the resolution of available abundance time series is low. On the other hand, the inclusion of trait data with either metric led both to an increase in EWS performance when faced with reduced sampling time-series resolutions and minimized any important differences between the two time-series metrics used. Thus, whenever possible, trait-based EWS will likely outperform abundance-based EWS in natural populations (but see (Baruah et al. 2019b)) and, when trait data is not available, abundance-based EWS derived from autocorrelation at-first-lag are more powerful and reliable indicators.

### 4.2 Effect of length of sampling time series on abundance-based and trait-based EWS

Reducing the length of the sampling time series in simulated data negatively affected the performance of abundance-based EWS. In our simulation study, having longer timeseries led to stronger EWS, particularly for slow and moderate levels of environmental forcing. In general, time series from slow and moderate forcing scenarios with lengths less than 5 generations returned weaker forecasts of population collapse with higher rates of false negatives (Fig. 2). The choice of fold or transcritical bifurcation models had little influence on predictability of population collapse, with the exception of the fast forcing scenario. When forcing was fast, transcritical models showed a shift in trend indicating the length of time series no longer had a strong effect on the predictability of collapse (Fig. 2 – dashed lines). In the context of a ciliate population such as the one used in the microcosms of this paper (*D. nasutum*) 10 generations corresponds to roughly 20 days. Twenty days’ worth of sampling might seem feasible to generate reliable forecasting for a ciliate population. However, for this same reliability in a population experiencing rapid forcing, 22 to 35 years of sampling would be required in the case of *Thynnus thynnus* (Atlantic bluefin tuna): an endangered species of high economic concern with a generation time between 2.2 to 3.5 years (Collette et al. 2011). For organisms with longer generation times such as *T. thynnus*, the required investment of resources and time becomes an important consideration if there is a desire to monitor populations using abundance-based EWS. In addition, we found that abundance time series require a minimum of 10 generations worth of data when forcing is fast and a minimum of 5 when forcing is moderate (Fig. 2). This finding suggests that in situations where forcing is more intense and, thus, populations are most at risk the requirement for good quality data is extended. This requirement of long time series is a clear shortcoming of abundance-based EWS for organisms with long generation times and populations experiencing rapid forcing.

In our microcosm study, the effect of time-series length on the Kendall’s tau metric was less clear than in the simulation. However, a general decreasing trend in forecasting strength was still observed in most forcing scenarios. In addition, abundance-based EWS rarely had the length of sampling required to make a confident forecast of collapse. It appears that regardless of improvements brought about by the increased length of the sampling time series, most abundance-based EWS did not provide a confident forecast even with the maximum length of sampling time series (between 10 and 12.5 generations). In our simulations we showed that abundance time series need a minimum of 10 generations of data to provide an accurate forecast. These microcosm results show that the required minimum length of sampling exceeds 10 generations in controlled laboratory settings. Thus, we can expect that in natural populations subjected to higher levels of stochasticity the minimum length of sampling might exceed even that which is required in the microcosms. However, in natural populations there is the possibility to measure phenotypic data and derive trait-based EWS which we have additionally assessed. The addition of phenotypic data with abundance data improved forecasting in the microcosms. Trait-based EWS, that included body size information alongside abundance-based EWS, offered a significant improvement over abundance-based EWS, providing a positive forecast of collapse for all time-series lengths tested.

Our modelling scenario of timeseries length and resolution was focused solely on abundance dynamics, and ignored trait dynamics. Abundance data are the most available, with databases such as Living Planet Index (LPI) or BioTime (Dornelas et al. 2018) offering an opportunity to analyze abundance time series data (>22,000) for future forecasts of population declines at a global scale. However, average time series length in the LPI databases is around 10.3 years and median length of 6 years. It is thus of foremost importance to understand whether short time series of different resolutions would affect the predictability of population decline before these statistical tools could be applied widely. Trait data, such as body size time series are less widely available. Hence, the main motivation of our work is to develop an understanding of the efficacy of abundance-based EWS in forecasting population decline when time series lengths are variable and are of varying resolutions.

Trait-based simulation of population dynamics could essentially be done using a quantitative genetic framework (see Baruah et al., 2019a). Infact, a recent simulation study, on comparing the strength of abundance-based and trait-based EWS, have implied the preferential use of trait-based EWS over abundance-based EWS. This particular study suggested that under certain ecological circumstances such, as high trait plasticity, and/or high reproductive rate, trait-based EWS outperformed abundance-based EWS in predicting population declines (Baruah et. al 2019b). Parallel to their simulation study, we show with experimental data from microcosms, that despite shorter timeseries lengths and/or low sampling resolutions, body-size based signals outperform abundance-based EWS. Whether a trait, such as body size, could be included among the suite of EWS will depend not only on the type of the trait but also on whether external environmental forcing affects the trait. If external environmental forcing does not affect body size it is expected that body size based EWS will fail in forecasting potential population declines. We thus expect traits that are correlated to fitness of an organism to be potential candidates to be included within the suite of trait based EWS.

That being said, body-size based signals are clearly potential candidates to be included alongside generic EWS of population collapse. Recent study on this aspect have highlighted the environmental, ecological, and evolutionary circumstances under which it is possible for phenotypic traits to shift before a potential population decline and thus act as a warning signal (Baruah et al. 2019a)). The strength of shifts in phenotypic trait such as body size, was dependent on how fast the environment changes, how plastic the trait is to changes in the external environment, and how high genetic variation was in the trait (Baruah et al. 2019). Infact, strength of trait-based EWS was directly dependent on the above ecological and evolutionary factors. For instance, it was suggested that, high levels of plasticity in the trait led to stronger trait-based EWS (Baruah et al. 2019b).

In our modelling scenarios, we chose a specific set of parameters to evaluate the performance of EWS in the face of myriad sampling lengths and resolutions. The choice of such a specific set of parameters that we used in our simulations were motivated through an iterative process that returned time series spanning a large range of forcing values while maintaining time series lengths. Changes in the parameter values of strength in forcing levels or growth rate did not significantly alter our simulation results (see appendix S1: Fig. S4-5). In fact, even when forcing levels and growth rate of the population was altered, we observe similar results: longer time series generally led to stronger forecasts of population collapse, while low resolution led to poor forecasts of population collapse with EWS.

Our findings on the effect of sampling length on the forecasting of population collapse by EWS suggest that the length of sampling required to calculate confident forecasts by abundance-based EWS are very long and likely impractical from a conservation and monitoring standpoint. On the other hand, trait-based EWS can give accurate and reliable forecasts with much less sampling data compared to their abundance-based counterparts.

## Conclusion

Our results stemming from simulations and a laboratory microcosm study make a case for the use of trait-based EWS over classical abundance-based signals based on their increased reliability and strength when faced with varying lengths and resolutions of sampling time series. We attempted here to evaluate these EWS with data which might more realistically resemble data gathered from field monitoring programs. However, true field data arising from monitoring programs are likely to be noisier than the data we have presented and analyzed here. Nonetheless, our results suggest that abundance-based EWS perform poorly when the length or resolution of abundance time series with regards to the process rate of the system is decreased. We found that including trait dynamics alongside abundance-based EWS to generate trait-based EWS leads to more reliable and confident forecasts of population collapse. We further found in our simulations that a length of ten generations and a resolution of a half generation are the minimum for deriving confident forecasts with abundance-based EWS. If trait-based EWS are used, this length and resolution requirement is much more relaxed. With these results in mind, we recommend considerations be made for the length and resolution of sampling time series required to accurately forecast populations in the design of monitoring programs. Further, we recommend the additional monitoring of phenotypic traits in populations, which have been shown to vary with increasing levels of environmental forcing such that trait-based EWS which have more forecasting power and reliability can be derived.

## Acknowledgments

The authors declare no conflict of interests. AA acknowledges support and funding from the University of Zurich BUSS program. The research was funded by ERC gran 337785 and Swiss National Science Foundation grant 31003A-182286 to AO and Forschungskredit Nr. FK-18-082 to GB.

