## Supplemental figures for "Effect of time-series length and resolution on abundance- and trait-based early warning signals of population declines"

**Appendix S1**

**Title: Effect of time-series length and resolution on abundance- and trait-based early warning signals of population declines**

**Authors:** Arkilanian A.A.[1], Clements C.F.[2,3], Ozgul A.[2], and Baruah G.[2]

*1 Department of Biology, McGill University, Montreal, Quebec, H3A 1B1, Canada.*

*2 Department of Evolutionary Biology and Environmental studies, University of Zurich, Winterthurerstrasse 30, 4055 Zurich*

*3 School of Biosciences, University of Melbourne, Parkville VIC 3052, Melbourne, Australia*

**APPENDIX S1**

**Table S1.** Rate of false negatives from abundance-based early warning signals (EWS) of population collapse in simulated populations led to collapse through harvesting at three different rates: weak, medium, strong. The rate of false negatives reported is the number of false negative signals returned divided by the number of true positive signals. Results are further split by four time-series resolutions tested (a value of 2 indicates sampling was performed every 2 generations) and by the metric used as an EWS.

| **Strength of forcing** | **Slow** | | | | **Moderate** | | | | **Fast** | | | |
|---|---|---|---|---|---|---|---|---|---|---|---|---|
| **Resolution** | *2* | *1* | *50* | *25* | *2* | *1* | *50* | *25* | *2* | *1* | *50* | *25* |
| **Sd** | 52.5 | 52.4 | 51.5 | 45.1 | 51.4 | 50.4 | 44.5 | 41.7 | 39.5 | 51 | 50.4 | 58.7 |
| **Ar(1)** | 46.5 | 46.1 | 49.7 | 46.6 | 46.4 | 47.8 | 48 | 42.6 | 52.5 | 46.5 | 48.2 | 60.7 |

**Table S2**: Adjusted $R_2$ and slope values for length of time series against Kendall's tau correlation coefficient for different levels of forcing (Slow, moderate and fast) and for two types of bifurcations (Fold and Transcritical).

| | | **Slow** | **Moderate** | **Fast** |
|---|---|---|---|---|
| **Fold** | ***Ar(1)*** | $R_2$ = 0.39<br>Slope = -0.01 | $R_2$ = 0.712<br>Slope = -0.09 | $R_2$ = 0.79<br>Slope = -0.04 |
| | ***Sd*** | $R_2$ = -0.009<br>Slope= -0.0003 | $R_2$ = 0.80<br>Slope = -0.07 | $R_2$ = 0.09<br>Slope = -0.03 |
| **Transcritical** | ***Ar(1)*** | $R_2$ = 0.433 | $R_2$ = 0.83 | $R_2$ = 0.39 |

| | | | | |
|---|---|---|---|---|
| | | Slope = -0.09 | Slope = -0.06 | Slope = 0.24 |
| | ***Sd*** | $R_2$ = 0.22<br><br>Slope = -0.075 | $R_2$ = 0.75<br><br>Slope = -0.1 | $R_2$ = 0.82<br><br>Slope = 0.80 |

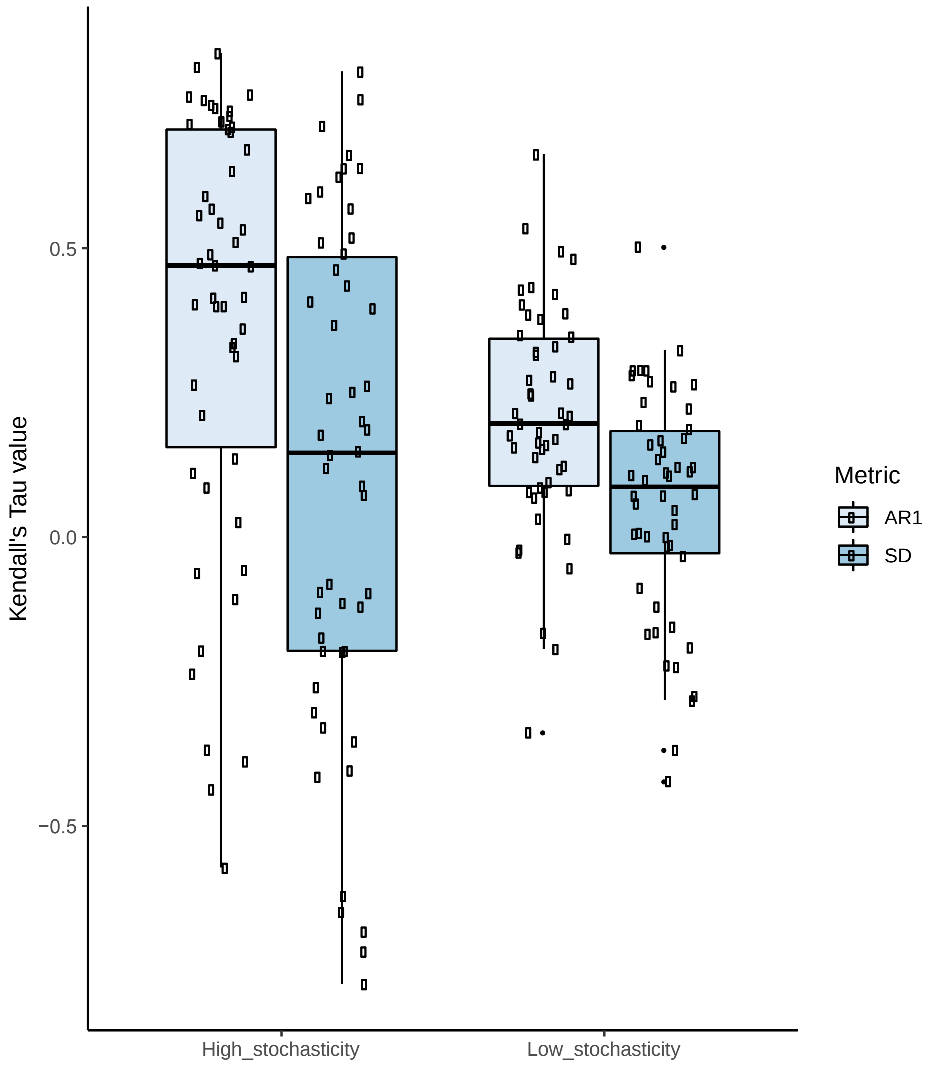


**Figure S1**: Performance (Kendall's tau value) of *ar(1)* and *sd* in forecasting population collapse for two different levels of environmental stochasticity (high and low) for 50 replicates of simulation collapse. Higher the value of Kendall's tau stronger is the early warning signal (*ar(1)* or *sd*) in forecasting population collapse.

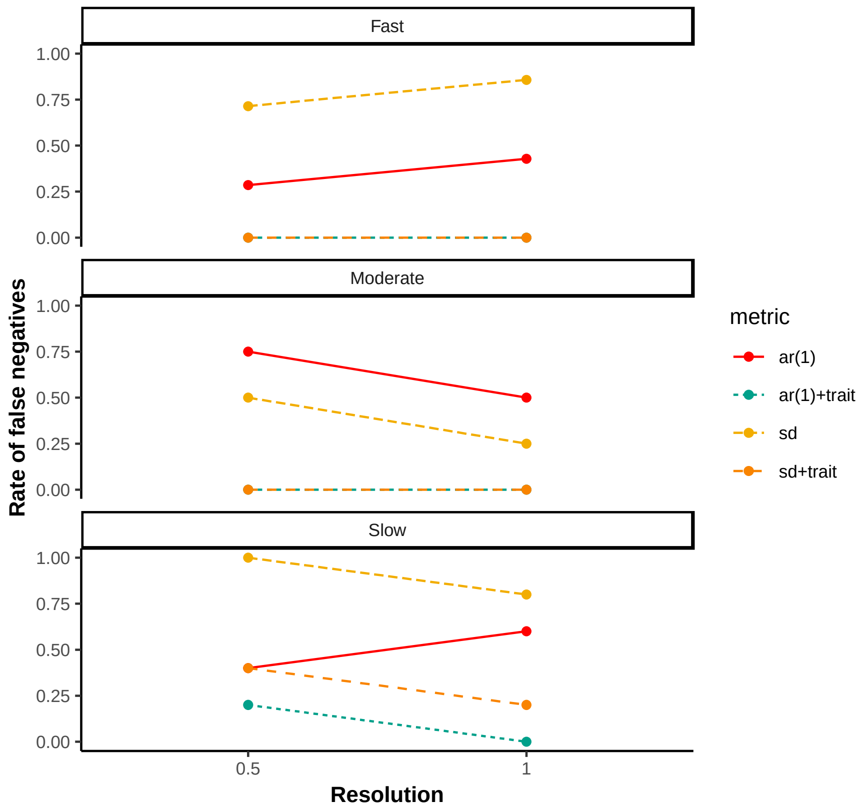


**Figure S2**: Rate of false negatives for the abundance (ar(1) , sd) and trait-based EWS (ar(1) +trait, sd+trait) for different time series resolution of the experimental data. Rate of false negatives are substantially lower for trait-based EWS in comparison to abundance-based EWS.

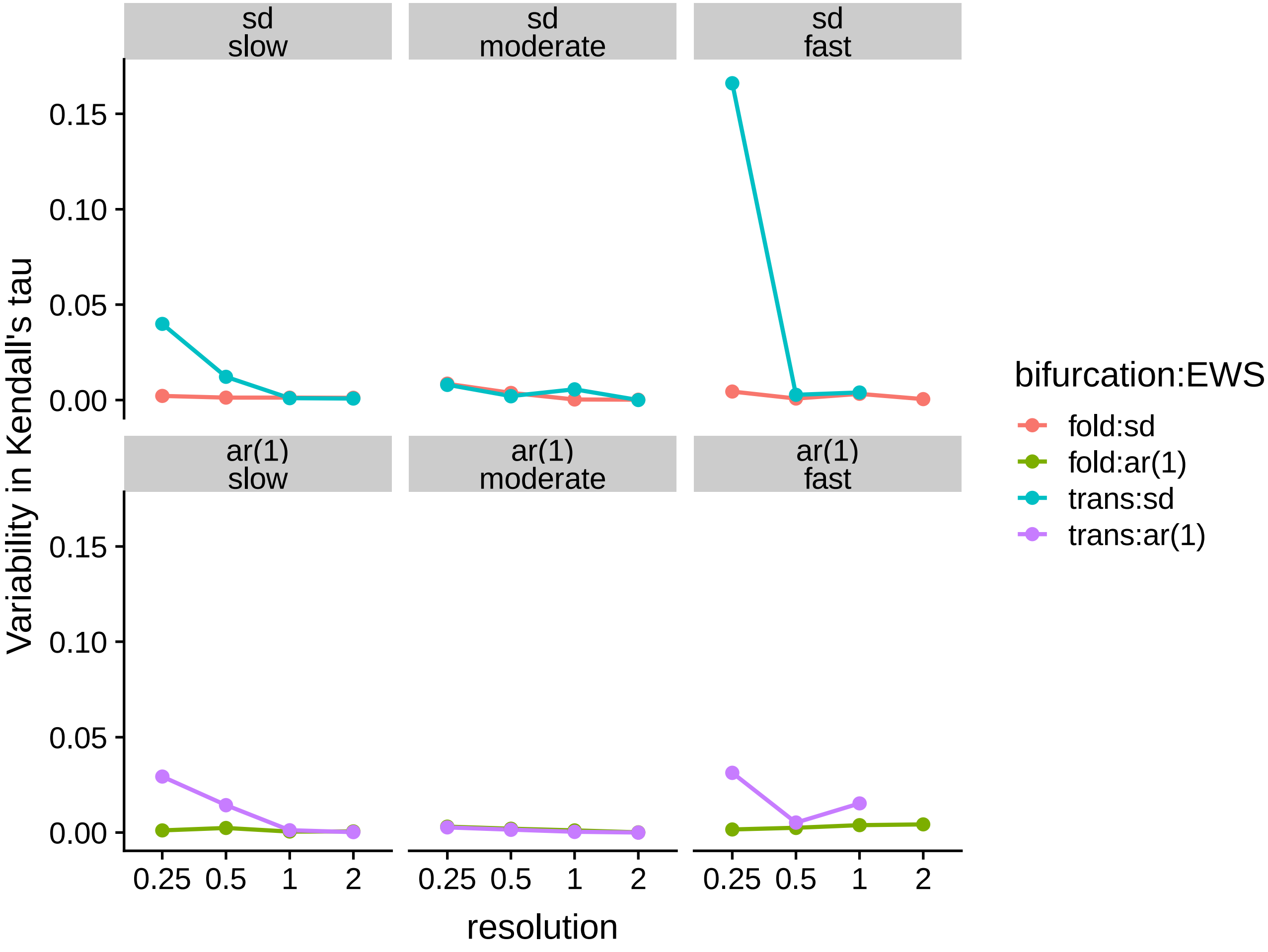


**Figure S3**: Variability (variance) in Kendall's tau correlation coefficient for different time series resolution for the simulated data. Top panel is for *sd* for different levels of environmental forcing (slow, moderate, fast) and the bottom panels are for *ar(1)* for slow, moderate and fast forcing. In general, variability in Kendall's tau decreased as the resolution decreased across the different levels of forcing both for *sd* and for *ar(1)*.

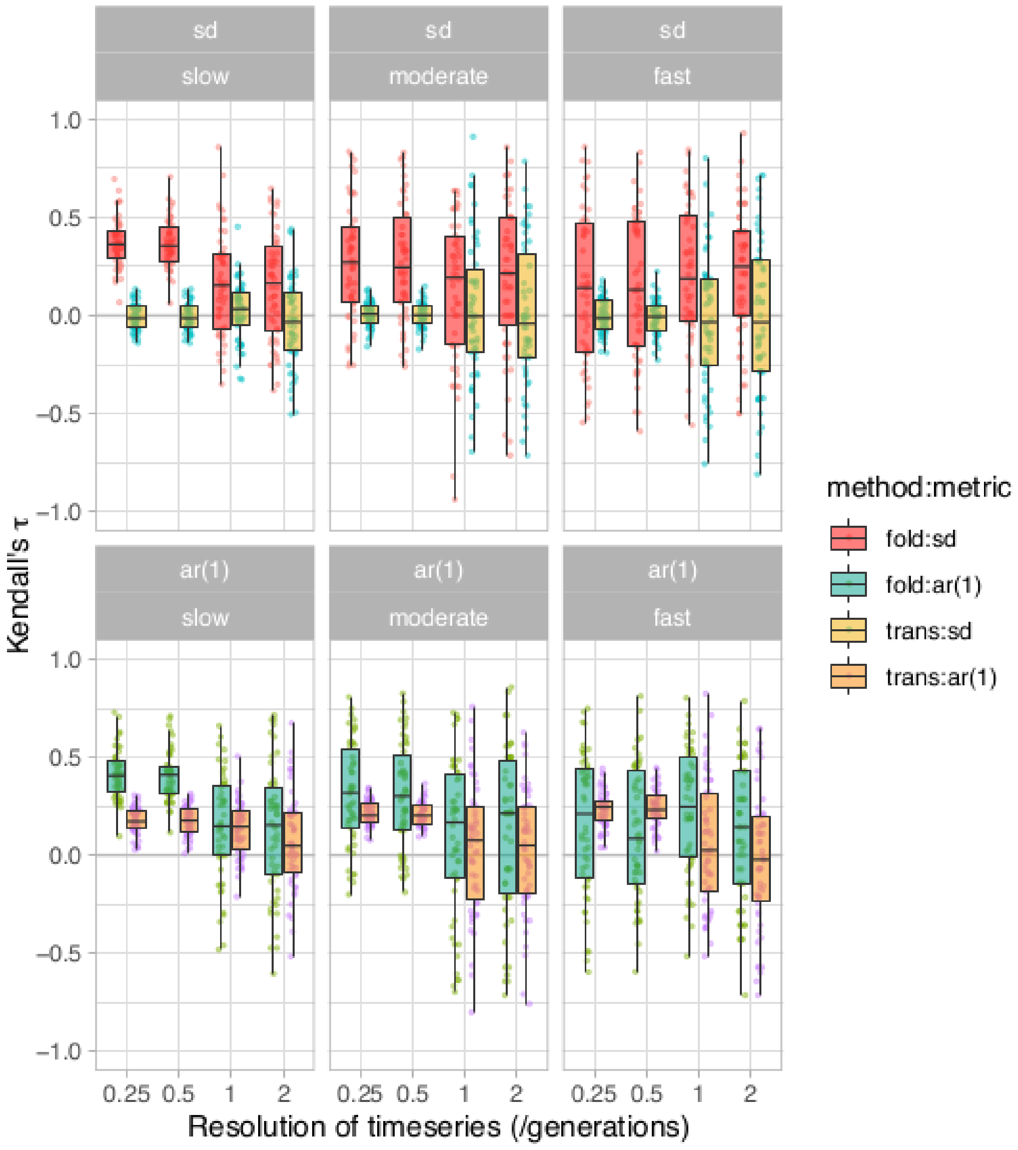


**Figure S4:** Boxplots of the strength of abundance-based early warning signals of population collapse across decreasing resolutions of sub-sampling in simulated populations subjected to collapse by harvesting, with different parameters used than the ones used for the main-text. The data is split into subplots based on the bifurcation model simulated (fold, transcritical) and forcing level (slow, medium, fast). Boxplots are further split by metric used (yellow: standard deviation, red: autocorrelation at-first-lag). Each box represents the median Kendall's tau value (shown on the y-axis) across replicate time series across three forcing level with lengths ranging from 1,000 to 40 time steps. The x-axis shows the resolution of sampling in numbers of generations such that a value of 2 indicates that sampling was performed every 2 generations and 0.25 denotes sampling every quarter generation. The parameters used for fold bifurcation are : $r = 1.2$ , K = 100, slow forcing rate = 0.3, medium forcing rate = 0.5, fast forcing rate = 0.7 . For transcritical bifurcation models the parameters used were : r =1.2, K= 100, slow forcing rate = 0.005, medium forcing rate = 0.007, and fast forcing rate = 0.009.

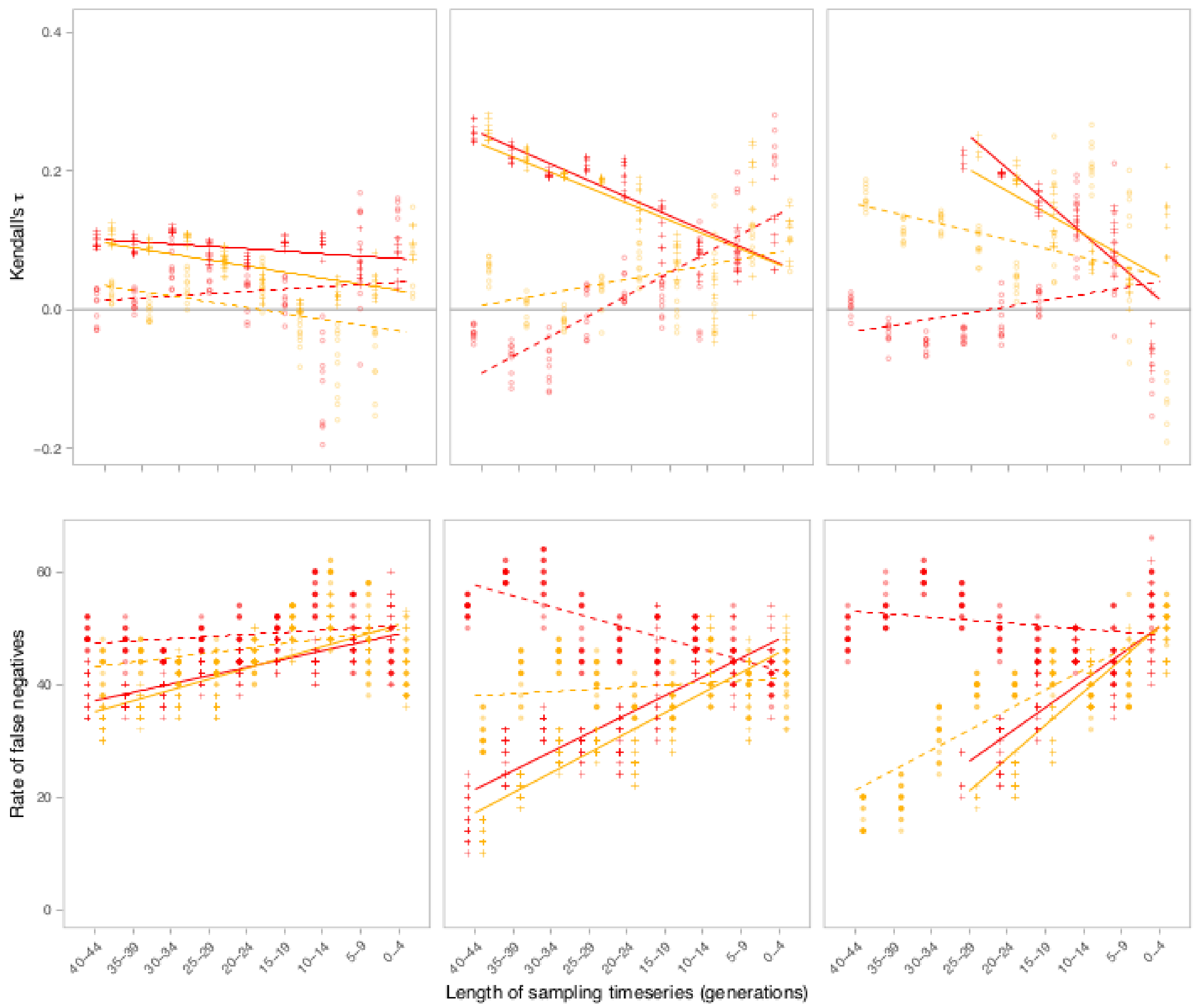


**Figure S5:** Performance of abundance-based early warning signals of population collapse across decreasing sampling time-series lengths in simulated populations subjected to collapse by harvesting for additional parameter values different from what was used for the main-text simulations. The data is split into subplots based on forcing intensity (slow, moderate, fast) and further split by metric used (yellow: standard deviation, red: autocorrelation at-lag-1) and bifurcation model simulated (solid: fold model, dashed: transcritical model). Each point represents (A) the mean Kendall's tau correlation coefficient or (B) rate of false negatives on the y-axis of 50 replicate simulations of population collapse. X-axis represents length of time series analyzed. The parameters used for fold bifurcation are : $r = 1.2$ ,  K = 100, slow forcing rate = 0.3, medium forcing rate = 0.5, fast forcing rate = 0.7 . For transcritical bifurcation models the parameters used were : r =1.2, K= 100, slow forcing rate = 0.005, medium forcing rate = 0.007, and fast forcing rate = 0.009.

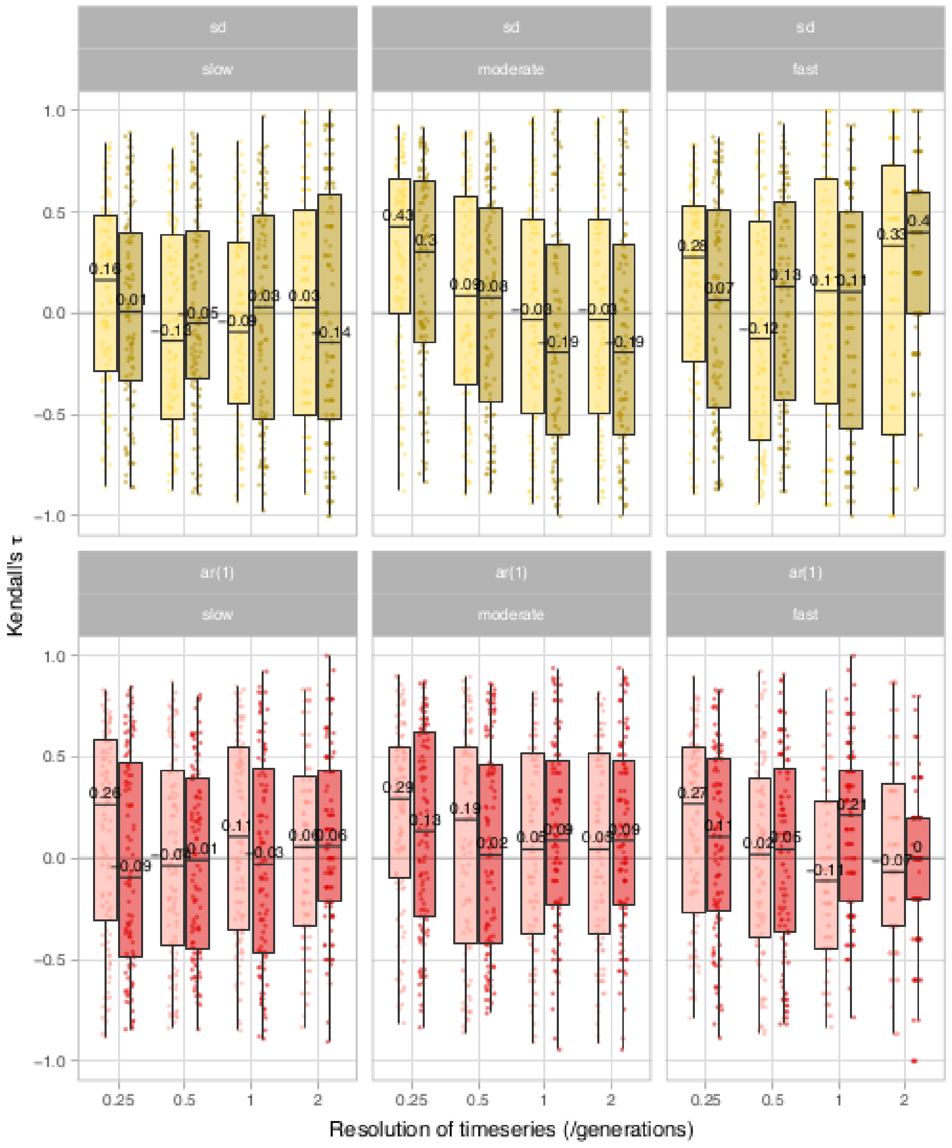


**Figure S6:** Boxplots of the strength of abundance-based early warning signals of population collapse across decreasing resolutions of sub-sampling in simulated populations subjected to collapse by harvesting, with median values written with each of the boxplots. This figure is the same as figure 1 of the main-text but with the median values noted. The data is split into subplots based on the bifurcation model simulated (fold, transcritical) and forcing level (slow, medium, fast). Boxplots are further split by metric used (yellow: standard deviation, red: autocorrelation at-first-lag). Each box represents the median Kendall's tau value (shown on the y-axis) across replicate time series across three forcing level with lengths ranging from 1,000 to 40 time steps. The x-axis shows the resolution of sampling in numbers of generations such that a value of 2 indicates that sampling was performed every 2 generations and 0.25 denotes sampling every quarter generation.
